# Inhibition of CELA1 Improves Septation in the Mouse Hyperoxia Model of Impaired Alveolar Development

**DOI:** 10.1101/2024.06.13.598911

**Authors:** Noah J. Smith, Rashika Joshi, Hitesh Desmukh, Jerilyn Gray, Andrea D. Edwards, Elham Shahreki, Brian M. Varisco

## Abstract

A key feature of bronchopulmonary dysplasia (BPD) is impaired alveolar septation. In later live, BPD survivors are more susceptible to childhood respiratory problems and have reduced respiratory function as adults. Chymotrypsin-like elastase 1 (CELA1) is a serine protease expressed in AT2 cells that mediates emphysema progression in adult mouse models. CELA1 binds and cleaves tropoelastin in response to strain. Its expression is developmentally regulated. Using the mouse hyperoxia model of impaired alveolar development we hypothesized a role for CELA1 in impaired alveolar development (IAD). In C57BL6 mouse pup lungs exposed to 80% oxygen for 14 days *Cela1* mRNA increased 1.9-fold (p<0.05) and protein 2.6-fold (p<0.01). Protein levels normalized after 14 days in room air. Analysis of an existing single cell mRNA-seq dataset showed *Cela1* mRNA in AT2 cells, alveolar macrophages and interstitial macrophages. The fraction of cells with Cela1 *mRNA* increased with hyperoxia. By flow cytometry the only *Cela1*-specific difference in immune cell populations was a 2-fold increase in lung eosinophils in room air (p<0.05). After 14 days of exposure to 80% oxygen *Cela1*^*-/-*^ mice had better alveolarization with an average mean linear intercept of 80 μm compared to 111μm (p<0.001). Treatment of hyperoxia-exposed pups with subcutaneous anti-Cela1 KF4 antibody offered similar protection compared to IgG (59 μm vs. 67 μm, p<0.001).Human BPD specimens demonstrated CELA1 in AT2 cells and myeloid cells. These data indicate that hyperoxia-induced increases in CELA1 are partially responsible for IAD and suggest a potential role in premature neonates exposed to high FiO_2_.

## Introduction

First described by Northway and colleagues in 1967, bronchopulmonary dysplasia (BPD) is a common sequelae of premature birth marked by failure of normal alveolar septation (1). Survival of prematurely born infants during this pre-surfactant replacement therapy era was largely limited to those with gestational age of 32 weeks or greater, and the lung histology of infants with “classic” BPD was marked by alveolar enlargement and fibrotic changes due to the barotrauma and volutrauma from the high ventilator pressures need to in these low compliance lungs (2). With the advent of surfactant replacement therapy, the gestational age of viability was reduced to ∼24 weeks. Additional advances have further reduced the age of viability to ∼22 weeks (3, 4). Surfactant replacement allowed for the use of lower ventilation pressures and earlier liberation from invasive mechanical ventilation, and new histological features were noted. This “new” BPD (now over 30-years old) was marked by failure of secondary alveolar septation and lacked the fibrotic changes of the “old” BPD. Surviving prematurely born infants with BPD are almost always liberated from invasive mechanical ventilation, but often require months or years of supplemental oxygen to maintain acceptable saturation levels. As both a treatment for and a cause of impaired alveolar septation, this presents a therapeutic conundrum for clinicians.

*Chymotrypsin-like Elastase 1* (*CELA1*) is a serine protease expressed by alveolar type 2 (AT2) cells (5) in a developmentally regulated manner (6) and is partially responsible the reduction of postnatal lung elastance. It binds to the non-crosslinked, hydrophobic domains of tropoelastin when exposed by strain and cleaves the molecules at these molecules. Although it is a digestive enzyme, it is universally conserved in placental mammals while other paralogues (*CELA2A, CELA2B, CELA3A, CELA3B*) have been variably lost suggesting a non-digestive role for *CELA1* (7). Its expression is increased with strain (8), and it is covalently bound and neutralized by α1-Antitrypsin (AAT) (7). When mechanical strain is applied to *ex vivo* lung sections, intrinsic elastase activity is increased in the vector of that strain (9), and CELA1-neutralizing antibody reduces this increase by ∼90% (5). *Cela1*-deficient mice are protected from emphysema in AAT deficiency (5, 10, 11), cigarette smoke-induced emphysema (5), and age-related alveolar simplification (5, 10). Interestingly and for unclear reasons, *Cela1*^*-/-*^ lungs have a higher percentage of leukocytes than wild type lungs (5, 10). A developed anti-CELA1 antibody inhibits CELA1 elastolytic activity and is similarly protective in these adult mouse models (5, 12). However, since Cela1 is responsible for ∼20% of the intrinsic elastin remodeling activity in the developing lung and *Cela1*^*-/-*^ mice have lower compliance lungs and slightly smaller alveoli than wild type mice (7), and because immune cells have an important role in alveolarization (13), it was unclear whether its ablation or inhibition would be beneficial in neonatal hyperoxia models of impaired alveolar development. We sought to test this hypothesis with particular attention to lung immune cell populations.

## Materials and Methods

### Regulatory Approvals

Animal use was approved by the Cincinnati Children’s Hospital Medical Center Institutional Animal Use and Care Committee (2020-0054). Human tissue was utilized under a waiver from the Cincinnati Children’s Hospital Medical Center IRB (2016-9641).

### Animal Use and Hyperoxia Model

All mice were on the C57BL/6 background, obtained from Jackson Laboratory, and provided water and autoclaved chow *ad libitum* in a barrier facility with 12-hour light-dark cycles. Female mice that had produced one previous litter were mated for one day, and matched pregnant wild type or *Cela1*^*-/-*^ pairs were allowed to deliver. Litters were culled to a maximum of 8 pups and half of each litter transferred with bedding to the matched dam. One litter was exposed to 80% oxygen and the other maintained in room air. Every 2 days the dams and half the cage bedding was switched. After fourteen days, pups were sacrificed.

### KF4 anti-CELA1 Antibody Administration

Mouse pups were administered 10, 20, and 35 μg of KF4 in PBS at a concentration of 1 mg/mL or mouse anti-Human IgG (Jackson Immunoresearch) subcutaneously between the shoulder blades based on a dose of 5 mg/kg (5) and an average mass of ∼2, 4, 7 mg on each of those days respectively (14).

### Tissue Processing and Histology

After anesthetization and sacrifice by exsanguination, the mouse pup thorax was opened trachea cannulated with a 22 gauge blunt needle and secured with silk suture, left bronchus ligated, left lung snap frozen, right inflated with at 30 cm H_2_O pressure using 4% paraformaldehyde in PBS, right lung fixed overnight, and lung lobes separated and paraffinized. Left lungs were homogenized with extraction of RNA using RNeasy columns (Qiagen) or processed for protein analysis. Right lung lobes were randomly oriented and embedded, sectioned, and stained using hematoxylin and eosin. Using the methods of Dunnill, *et al* (15), mean linear intercepts were determined in 5 fields per lobe.

### Lung Cell Isolation

Animals were sacrificed as above but inflated and incubated with Dispase (BD Biosciences) for 45 minutes at 37°C. Lungs were transferred to a 100 cm^2^ petri dish with MEM and 10% FBS and minced removing the trachea and bronchi. The suspension was passed through a 70-μm strainer, pelleted, and re-suspended red cell lysis buffer (BD Biosciences) for flow cytometry.

### Flow Cytometry

One million cells from each lung were incubated (4°C, 30 minutes) with anti–mouse CD16/CD32 (BD Biosciences, 553142) to block Fc receptors. The cells were reincubated (10 minutes, room temperature) with cell-surface marker antibodies (all diluted 1:100, Table 1). For intracellular staining, cells were washed and fixed (room temperature, 30 minutes) with the True-Nuclear Transcription Factor Buffer set fixative (BioLegend) and permeabilized (4°C, overnight) using 1× Permeabilization Buffer according to the manufacturer’s instructions. Cells were stained with intracellular antibodies (diluted 1:100, Table 1) and then washed twice and resuspended in flow cytometry buffer. Data were acquired with an Aurora (Cytek) and analyzed with FlowJo (Tree Star).

### Bioinformatic Analysis of Single Cell Data

Count matrices from the publicly available dataset GSE151974 (16) was downloaded and re-analyzed and data visualized using Seurat (17), sctransform (18), and pheatmap (19). After filtering by feature counts and percent mitochondrial gene reads, cells were clustered and groups defined using gene sets derived from UNCURL (20) in Table 2.

### Statistical Analysis

Using R version 4.0.2 (21), the following packages were used for statistical comparisons and graphics generation: ggpubr (22), ggplotify (23), and rstatix (24). For parametric data, Welch’s t-test and 2-way ANOVA with Holm-Šidák *post hoc* test were used, and for nonparametric data, Wilcoxon’s rank-sum and Kruskal-Wallis with Dunn’s *post hoc* test were used. All comparisons were 2-tailed. Parametric data are displayed as line-and-whisker plots, with the center line representing the mean and whiskers standard deviation. Nonparametric data are displayed as box-and-whisker plots, with the center line representing the median value, boxes representing the 25th to 75th percentile range, and whiskers representing the 5th to 95th percentile range. For both plot types, dots represent individual data points. For all analyses, P values of less than 0.05 were considered significant.

## Data Availability Statement

The scripts used for analysis and raw data are publicly available at 10.6084/m9.figshare.26035171.

## Results

### *Cela1* Expression in AT2 Cells and Macrophages Increases with Hyperoxia

We previously reported that *Cela1* is specifically expressed in mouse distal lung epithelial /alveolar type 2 cells (AT2) cells during lung development (6). By both PCR (Figure 1A) and Western Blot (Figure 1B), *Cela1* was increased in the lungs of mice exposed to hyperoxia. Analysis of a previously published single cell mRNA dataset in which C57BL6 mouse pups were exposed to 85% oxygen beginning at Postnatal Day 0 (PND0) and lungs collected at PND3, PND7, and PND14 identified that *Cela1* was expressed in both AT2 cells and myeloid cells (Figure 1C&D). In the AT2 cell cluster, there was a slight increase in the percentage of cells containing *Cela1* mRNA in hyperoxia after both 3, 7, and 14 days of hyperoxia exposure (Figure 1E-I). Among myeloid cells, *Cela1* was expressed mostly in alveolar and interstitial macrophages (Figure 1J&K). There was a more marked increase in the fraction of cells containing *Cela1* mRNA in alveolar and interstitial macrophages (Figure 1L-N). Taken together, unlike what we noted in adult mouse lungs (5, 7, 10), in the developing postnatal mouse lung, *Cela1* is expressed in AT2 cells, alveolar macrophages, and interstitial macrophages, and its expression increases in response to hyperoxia.

**Figure 1:**
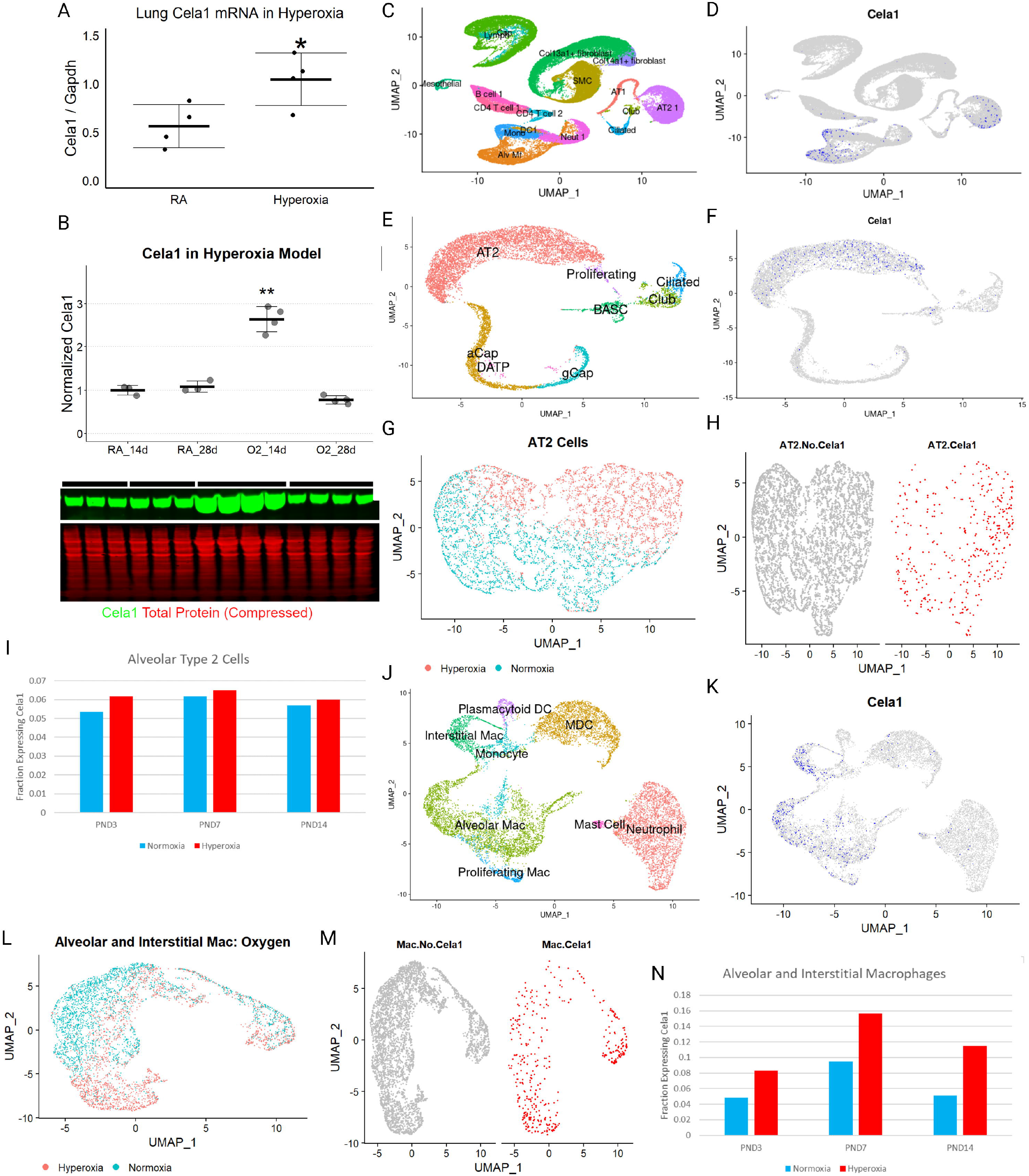
Hyperoxia Increases Cela1 Expression in AT2 Cells and Lung Macrophages. (A) After 14 days of exposure to 80% oxygen (hyperoxia), there is a doubling of lung *Cela1* mRNA in C56BL6 mouse pups compared to room air (RA). ^*^ p<0.05 by Student’s t-test. (B) Western Blot of the lungs of mouse pups exposed to 14 days of 80% oxygen (hyperoxia) and then 14 days of room air (RA). Cela1 protein levels are significantly higher after 14 days of hyperoxia and return to baseline after returning to room air. Compressed Western Blot displaying Cela1 protein (green) and total protein (red).. The bars above the blots represent three room air and four hyperoxia samples per time point and correspond to the plot labels. ANOVA p<0.01 and ^**^ indicates p<0.01 by holm *sidak post* hoc test of the 14-day hyperoxia group compared to all other groups. (C) Umap of a previously published single cell mRNA-Seq dataset of C57BL6 mouse pups exposed to 85% hyperoxia and assessed at PND3, PND7, and PND14. Cell clusters are annotated based on markers described in the supplement. (D) Feature plot of Cela1 mRNA-containing cells showing that the epithelial cell and myeloid cell clusters had contained Cela1-expressing cells. (E) Re-clustering of the epithelial clusters yielded the expected cell types. (F) *Cela1* mRNA was present in the AT2 cell cluster. (G) Among the AT2 cells, cells from hyperoxia-exposed mice largely separated along the second principal component and were unevenly distributed between the two major sub-types of AT2 cells. (H) There were slightly more Cela1-expressing AT2 cells in the cluster that had more hyperoxia-exposed AT2 cells. (I) Bar chart of relative fraction of AT2 cells containing Cela1 mRNA at PND3, PND7, and PND14. (J) Re-clustering of the myeloid cell clusters identified the expected cell types. (K) Cela1 mRNA was present in both alveolar macrophages and interstitial macrophages. (L) These two cell types also had groupings within each cluster based on exposure to hyperoxia. (M) Both the alveolar macrophage (left) and interstitial macrophage (right) clusters had increased Cela1-expressing cells in hyperoxia-exposed cells. (N) The relative fractions of alveolar and interstitial macrophages expressing *Cela1* were increased at all three time points.

### *Cela1*-deficient Mice Are Protected from Impaired Alveolar Development in the Hyperoxia Model

Wild-type and *Cela1*^*-/-*^ mice were exposed to 80% hyperoxia for 14 days using a time-matched, paired breeding strategy. After 14 days of hyperoxia exposure, *Cela1*^*-/-*^ mice exposed to hyperoxia had an average mean linear intercept (MLI) of 80 μm compared to 111 μm in wild type mice. *Cela1*^*-/-*^ hyperoxia mice had 63% protection compared to wild type mice exposed to room air. By two-way ANOVA the MLI effects of both a *Cela1* knock-out genotype (p=0.00014) and hyperoxia (p=2.7×10^-11^) were significant (Figure 2A-E). These differences in alveolar structure persisted after bringing mice into room air for 14 days with 51% protection with a *Cela1* knock-out genotype and hyperoxia effects of p=0.00026 and p=1.3×10^-11^ respectively (Figure 2F). In addition to re-confirming the previously described smaller alveolar size in *Cela1*^*-/-*^ mice (7), these data show partial protection of *Cela1*-deficient mice in the hyperoxia model of impaired alveolar development.

**Figure 2:**
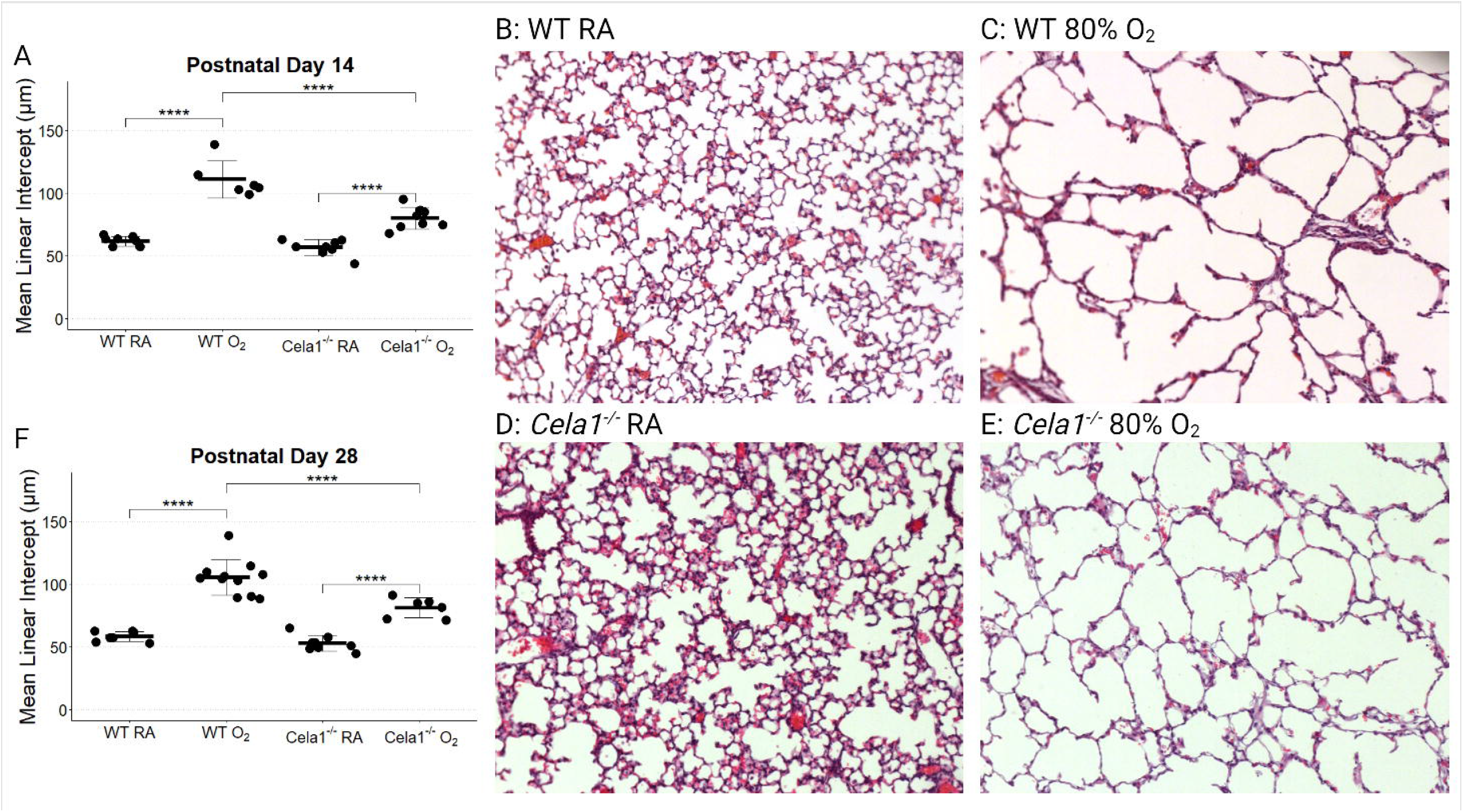
*Cela1* Deficiency Preserves Alveolar Structure in the Hyperoxia Model. (A) Hyperoxia caused a significant increase in airspace size in both wild type and *Cela1*^*-/-*^ mice, but this difference was less in *Cela1*^*-/-*^ mice. WT room air, WT hyperoxia, *Cela1*^*-/-*^ room air, and *Cela1*^*-/-*^ hyperoxia groups had 8, 6, 8, and 8 mice respectively. (B) Representative 10X photomicrographs of PND14 lung of wild type mice raised in room air, (C) wild type mice raised in 80% oxygen, (D) *Cela1*^*-/-*^ mice raised in room air, and (E) *Cela1*^*-/-*^ mice raised in 80% oxygen. (F) After returning hyperoxia mice to room air for an additional 14 days, the differences in MLI persisted. *Cela1*^*-/-*^ mice. WT room air, WT hyperoxia, *Cela1*^*-/-*^ room air, and *Cela1*^*-/-*^ hyperoxia groups had 7, 11, 8, and 6 mice respectively. ^****^p<0.0001 by tukey *post hoc* test after two-way ANOVA p<0.05.

### Few *Cela1*-mediated Differences in Lung Immune Cell Populations in Hyperoxia

Since *Cela1* expression increased in alveolar macrophages and interstitial macrophages in response to hyperoxia, we wondered whether Cela1-deficiency could have some impact on lung immune cell populations. We performed flow cytometry on lung cells from wild type and *Cela1*^*-/-*^ mouse pups after 14 days of exposure to 80% oxygen or room air (Figure 3A). As a fraction of total cells, exposure to hyperoxia induced a non-significant increase in neutrophils and a significant increase in Th2 lymphocytes, innate lymphoid cells, interstitial macrophages, and eosinophils. Hyperoxia induced a significant reduction in alveolar macrophages. The only cell population with differential abundance between wild type and *Cela1*^*-/-*^ was eosinophils with Cela1^-/-^ mice having more eosinophils in room air (0.15% vs 0.07%, p<0.05, Figure 3B). Overall, these data demonstrate significant differences in immune cell populations after exposure to hyperoxia and that the absence of *Cela1* has little impact on these.

**Figure 3:**
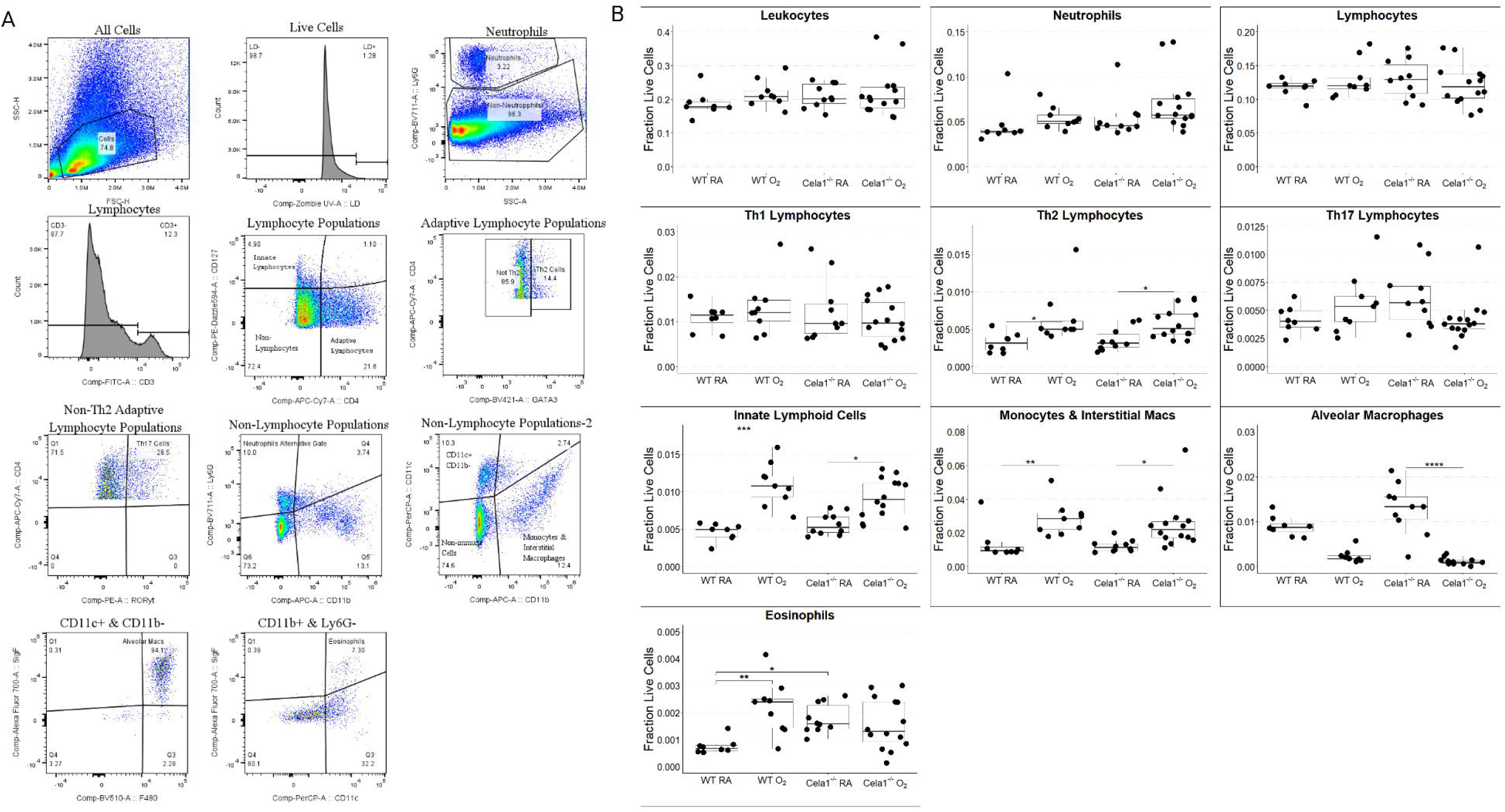
Immune Cell Quantification in Hyperoxia Model. (A) Cell Gating Strategy. Cells singlets were gated by forward scatter and side scatter. Dead cells were excluded. Neutrophils were gated by Ly6G, and lymphocytes were gated by CD3 in the Ly6G-negative population. Innate lymphoid cells were gated by CD127 and adaptive lymphocytes by CD4. CD4-positive cells were gated as Th2 on GATA3, and non-Th2 cells as Th17 by RORγt-positive and Th1 by RORγt-negative. In non-lymphocytes, additional neutrophils were gated on Ly6G at a lower threshold than previous, interstitial macrophages and monocytes were gated based on CD11b, and alveolar macrophages were gated on CD11c, SigF, and F4/80. The sum of the two neutrophil populations and sum of each major cell population were used for determination of fraction of neutrophils and leukocytes, respectively. (B) The lungs of 8, 9, 10, and 14 Wild type room air (WT RA), WT hyperoxia (O_2_), *Cela1*^*-/*-^ RA, and *Cela1*^*-/-*^ O2 exposed mice were analyzed. Hyperoxia increased the fraction of Th2 lymphocytes, innate lymphoid cells, monocytes, and interstitial macrophages and decreased the fraction of alveolar macrophages. Hyperoxia increased the fraction of eosinophils in wild type mice, but in *Cela1*^*-/-*^ mice levels were comparable to wild type hyperoxia and knockout hyperoxia lungs. ^*^p<0.05, ^**^p<0.01, ^***^p<0.001, ^****^p<0.0001 by Dunn’s *post hoc* test after Wilcoxon rank-sum p<0.05.

### KF4 anti-Cela1 Antibody Protects Against Impaired Alveolar Development

Since there was evidence that increased Cela1 levels contributed to hyperoxia-induced impaired alveolarization, we treated neonatal mouse pups at 3, 7, and 10 days with subcutaneous anti-Cela1 KF4 antibody. This antibody has been previously described to bind to the histidine of the Cela1 catalytic triad (5). Similarly dosed mouse anti-Human IgG was used as a control. KF4 treatment had similar protection against impaired alveolarization as was seen in *Cela1*^*-/-*^ animals (Figure 4A-E). These data indicates that at least in this mouse model of BPD, neutralizing hyperoxia-induced increases in Cela1 can preserve alveolar microstructure.

**Figure 4:**
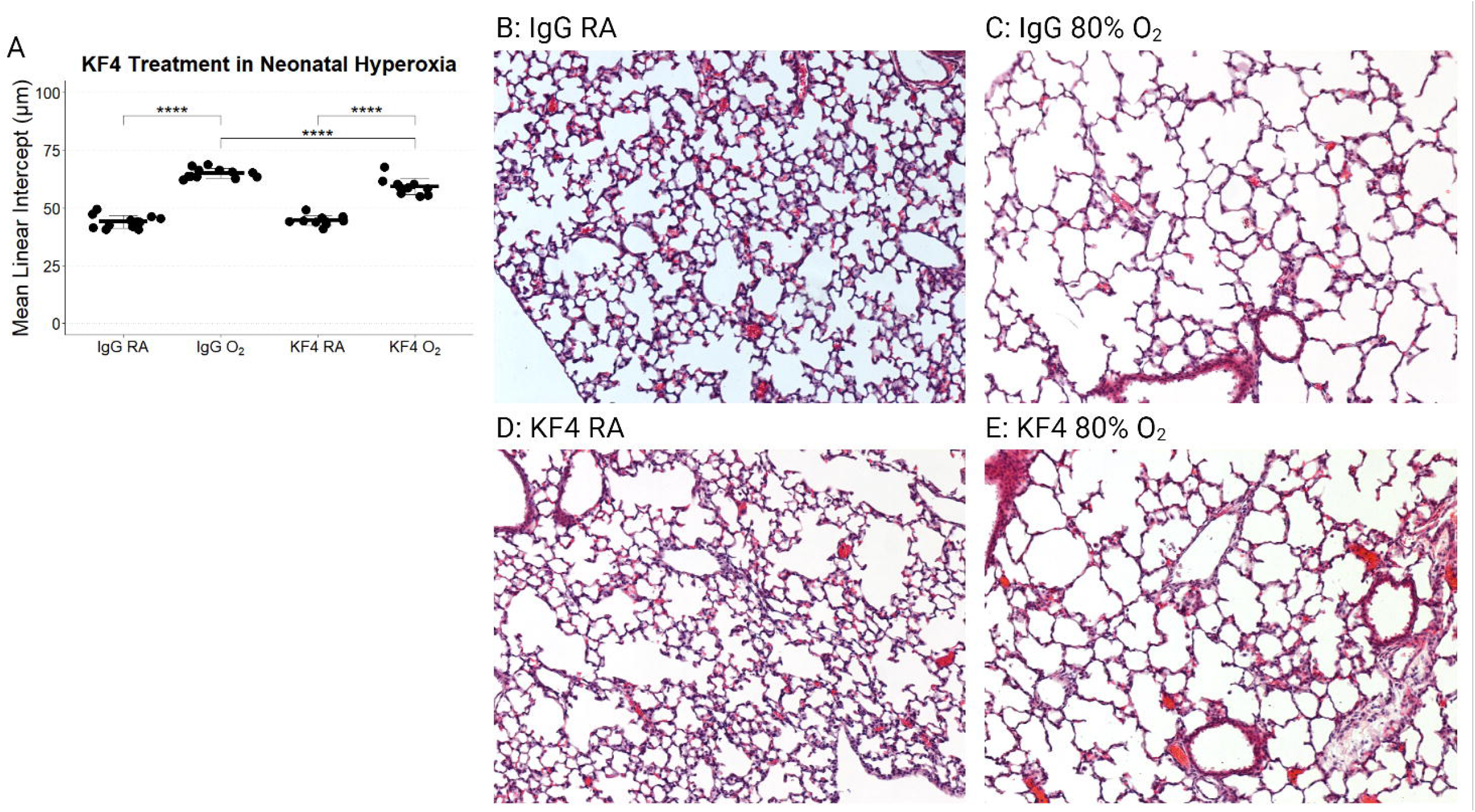
KF4 anti-CELA1 Antibody Preserves Alveolar Structure in the Hyperoxia Model. (A) Both IgG and KF4-treated mice had a significant increase in alveolar size when exposed to hyperoxia. However, KF4-treated mice had less alveolar simplification than IgG-treated mice. IgG room air, IgG hyperoxia, KF4 room air, and KF4 hyperoxia groups had 12, 12, 10, and 11 mice respectively. (B) Representative 10X photomicrographs of PND14 lung of IgG-treated mice raised in room air, (C) IgG-treated mice raised in 80% oxygen, (D) KF4-treated mice raised in room air, and (E) KF4-treated mice raised in 80% oxygen. ^****^p<0.0001 by tukey *post hoc* test after two-way ANOVA p<0.05.

### CELA1 is Present in AT2 cells in the Lungs of Infants with Bronchopulmonary Dysplasia

There are very few lung specimens available of infants with BPD. Immunofluoresecent and proximity ligation *in situ* hybridization imaging of term infant lung identified few CELA1-containing cells (Figure 5A). The lung of an infant with BPD demonstrated more expressing cells and co-localization of *CELA1* mRNA and protein (Figure 5B). Immunohistochemistry for CELA1 identified few CELA1-containing cells in term infant lung (Figure 5C), and in BPD lung several cells with macrophage morphology and location had vesicular staining for CELA1 (Figure 5D).

**Figure 5:**
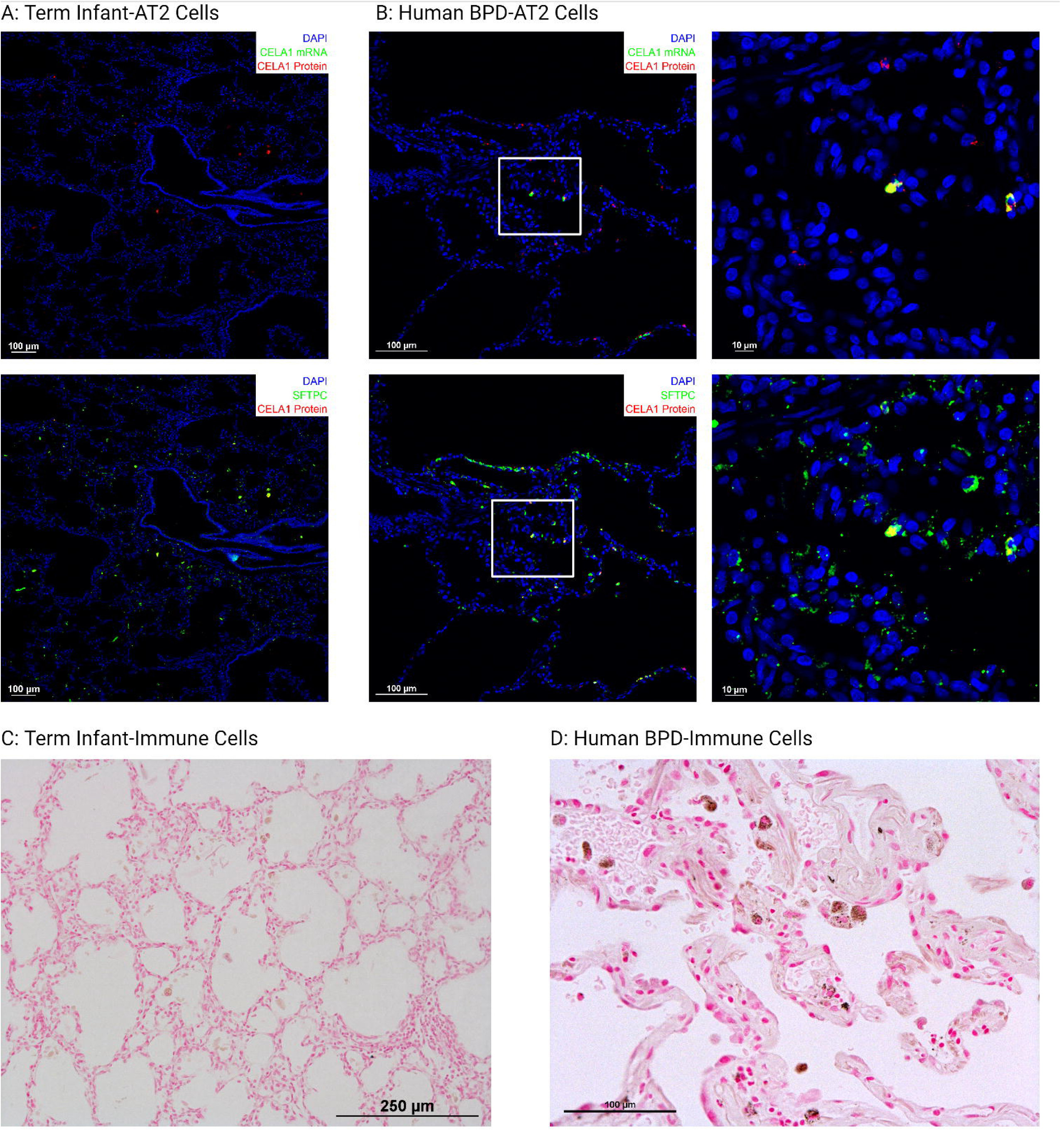
CELA1 in Human BPD Lung. (A) Tissues from an 8 month-old infant born at 25 weeks gestation with BPD demonstrated CELA1-mRNA (green) and CELA1 protein (red) containing cells. In a separate channel, CELA1 protein was within pro-Surfactant Protein C (Green) containing cells. (B) A separate section showed CELA1 protein in immune cells labeled with CD45 (white). (C) Tissue from a 1 week-old infant born at 40 weeks gestation without known lung disease showed rare CELA1-protein expressing AT2 cells.

## Discussion

This study is the first to demonstrate a role for CELA1 in hyperoxia-induced alveolar simplification during a developmentally crucial time window. Unlike adult lung in which CELA1 expression is largely limited to AT2 cells, we found that in the developing lung, *Cela1* was also expressed in alveolar and interstitial macrophages. Hyperoxia increased expression in all three cell types and expression returned to baseline after hyperoxia exposure ceased. Limited human lung data showed that similar expression patterns may be present in premature infant lungs. Thus, the KF4 anti-CELA1 antibody could represent a novel therapy to attenuate hyperoxia-induced alveolar simplification.

It is unlikely that CELA1 mediates hyperoxia-induced alveolar damage directly. CELA1 is a serine protease that binds and cleaves non-crosslinked hydrophobic tropoelastin residues under conditions of strain (7, 8). KF4 directly binds to the histidine of the CELA1 catalytic triad (5) presumably preventing this binding. The elastin architecture of mice exposed to hyperoxia (25) and infants with BPD (26) is disordered, and both elastin and collagen networks are crucial for the even distribution of lung strain (27). If CELA1 remodels lung elastin in the developing lung in a similar manner to the adult lung (10), then excessive strain in some lung regions could lead to aberrant remodeling of these regions. This would represent developmentally inappropriate or excessive remodeling as the physiologic role for *Cela1* in the developing lung is to reduce postnatal elastance (6, 7). In room air experiments, *Cela1*^*-/-*^ lung had slightly smaller MLI than wild type lung as was previously reported, although the significance of this was lost after accounting for repeated measures.

It is unclear which cell type is responsible for CELA1-mediatied remodeling. CELA1 secreted from both AT2 cells and interstitial macrophages would have access to elastin fibers in the interstitial space. Both of these cell types are located in regions in which strain could be sensed, and so having effector mechanisms to reduce this strain makes sense. CELA1 secreted from alveolar macrophages are not so located and would not have access to the interstitial compartment, so the mechanism by which CELA1 secreted by these cells would act is uncertain. Since α1-antitrypsin neutralizes CELA1, a regulatory mechanism is potentially present within the interstitial space but not the alveolar space.

Several weaknesses in the current study should be noted. First, due to the paucity of available specimens, we only evaluated one BPD and one control specimen by histology. The ages of these specimens (8 months and 1 week old respectively) were discrepant, and we cannot know whether the infant with BPD was receiving supplemental oxygen at the time of death. While additional specimens could be gathered, it is unlikely that meaningful, quantitative data could be generated from autopsy specimen. We did not quantify the extent to which KF4 reduced lung elastolytic activity as we did following tracheal administration of porcine pancreatic elastase in adult mice (5, 12). We did not perform dosing experiments as we did for intraperitoneal administration of KF4, so it may be that higher or lower doses of KF4 would have greater efficacy or less potential toxicity. Notably, doses of 25 mg/kg intraperitoneal KF4 or IgG had minimal toxicity in adult mice (12). In flow cytometry experiments, we used cell-specific markers but not the general marker CD45 to identify immune cells. It may be that using the sum of all positive cells to define leukocytes omitted an important population explaining why we saw a trend but not significant increase in the fraction to total cells that were immune cells as we have in previous studies (5, 12).

There are currently no therapies that mitigate the effects of hyperoxia on impaired alveolar development. Since this is a key feature of BPD, future work will require an evaluation of CELA1 levels in the tracheal aspirate fluid of prematurely born infants exposed to hyperoxia. Since previously published microarray data in chronic obstructive pulmonary disease detected higher CELA1 levels in the sputum of subjects with advanced stage COPD (28, 29), it may be feasible to directly measure CELA1 mRNA in the tracheal aspirates of intubated prematurely born infants to test for an association with inspired oxygen levels.

## Acknowledgements

We would like to thank the LungMAP Human Tissue Core at the University of Rochester Medical Center for providing human lung specimens and the Cincinnati Children’s Hospital Flow Cytometry and Confocal Imaging Cores.

## Abbreviations

AT2 Cell: Alveolar Type 2 Cell
BPD: Bronchopulmonary Dysplasia
CELA1: Chymotrypsin-like Elastase 1
FiO_2_: Fraction inspired O_2_
IAD: Impaired Alveolar Development
PND: Postnatal Day

## Declarations

Cincinnati Children’s Hospital and BMV hold patents (US 63009134 and20200413) for KF4-based therapies for the treatment of AAT-deficient and non–AAT-deficient emphysema (case 2016-0706).

## Notes

### Competing Interest Statement

Cincinnati Childrens Hospital and BMV hold patents (US 63009134 and 20200413) for KF4 based therapies for the treatment of AAT-deficient and non AAT deficient emphysema (case 2016-0706).

